# Evolutionary Insights from a Large-scale Survey of Population-genomic Variation

**DOI:** 10.1101/2023.05.03.539276

**Authors:** Zhiqiang Ye, Wen Wei, Michael Pfrender, Michael Lynch

**Author notes:** Co-first authors (Zhiqiang Ye, Wen Wei).

## Abstract

Results from data on *>* 1000 haplotypes distributed over a nine-year period from a natural population of the microcrustacean *Daphnia pulex* reveal evolutionary-genomic features at a refined scale, including key population-genetic properties that are obscured in studies with smaller sample sizes. Background selection, resulting from the recurrent introduction of deleterious alleles, appears to strongly influence the dynamics of neutral alleles, inducing indirect negative selection on rare variants and positive selection on common variants. Fluctuating selection increases the persistence of nonsynonymous alleles with intermediate frequencies, while reducing standing levels of variation at linked silent sites. Combined with the results from an equally large metapopulation survey of the study species, regions of gene structure that are under strong purifying selection and classes of genes that are under strong positive selection in this key species can be confidently identified. Most notable among rapidly evolving *Daphnia* genes are those associated with ribosomes, mitochondrial functions, sensory systems, and lifespan determination.

---

With the advent of genomic sequencing, substantial progress has been made in understanding the genetic structure of populations at the genome-wide level, although large-scale studies of this sort are still confined to a few species. From single-generation samples alone, impressive insights have have been gained from studies on model organisms such as *Drosophila* (e.g., Lack et al. 2015; Everett et al. 2020; Kapun et al. 2021) and humans (Lachance and Tishkoff 2013; Byrska-Bishop et al. 2022; Fatumo et al. 2022). However, even for the most commonly studied model species, uncertainty remains about fundamental features such as genotypic-frequency agreement with Hardy-Weinberg expectations, the stability of allele frequencies across generations, and the roles played by drift vs. purifying and/or positive selection on various gene categories and gene regions.

This study reports on a 9-year (*∼* 40-generation) survey of population-genomic data for the aquatic microcrustacean *Daphnia pulex.* With each sample being derived from newly hatched, sexually produced progeny, and annual intervening periods between bouts of sex being *∼* 5 generations of clonal reproduction, the genetics of the study population are quite similar to those found in other purely sexual species such as *Drosophila* and humans, including average rates of recombination (Lynch et al. 2017). Early studies on *Daphnia* populations based on allozymes suggested changes in allele frequencies across years far greater than expected by random genetic drift (Lynch 1987), although the causal mechanisms remained unclear, and by necessity the prior work was focused on a very small number of genes with high-frequency alleles. Uncertainties with the earlier work are clarified by the temporal series of data for the study population, which revealed a number of statistical features with respect to the magnitude and spatial patterning of selection across the genome (Lynch et al. 2023), and provides the backdrop for the more extensive gene-specific analyses in the current paper.

As it involves the analysis of *∼* 700 diploid individuals, this study represents one of the most thorough population-genomics analysis of any organism. The justification for such sampling effort is not simply that larger sample sizes yield smaller standard errors of parameter estimates, but that key population-genetic features can be completely obscured with single-point samples involving 100 or fewer individuals (Cvijovíc et al. 2018; Buffalo and Coop 2019; Lynch and Ho 2020). Supplementing the temporal analysis of Lynch et al. (2023), the analyses presented herein provide further insight into the basic population-genetic features of *D. pulex,* including the site-frequency spectra of rare alleles, the magnitude of linkage disequilibrium, and the genome-wide pattern of selection operating on various functional categories of genes. We also identify outlier genes under extreme forms of purifying vs. positive selection, providing insight into the key targets of selection in the study species. Throughout, we draw comparisons from a recent study of single-year samples for ten populations of the study population, referred to below as the metapopulation study (Maruki et al. 2022). Together, these two large studies set the stage for future work on the evolutionary and ecological genomics of key gene functions.

## RESULTS

The study population occupies an ephemeral, natural woodland pond in a Nature Conservancy tract from Portland Arch, Fountain County, Indiana. The following analyses rely on a temporal series of genome-sequence data from annual population samples consisting of 72 to 92 diploid individuals (Supplemental Table S1 in Lynch et al. 2023). Each sample was obtained shortly after resting-egg hatch-out (typically mid-March to mid-April) and hence represents first-generation sexually produced offspring emerging prior to the operation of selective events during the subsequent *∼* 3 to 5 generations of clonal reproduction. Generally, the population enters a phase of sexual reproduction and resting egg production by June, as the pond dries up and remains so until the following spring.

### Patterns of standing variation

The overall coverage and numbers of genotypes sampled per year is sufficient for essentially all nucleotide sites with annual estimated minor-allele frequencies (MAFs) *>* 0.05 to be deemed significant (i.e., not due to sequencing error) in any particular sample (Figure 1). Between *∼* 5 and 23% of nucleotide sites with MAFs greater than this cutoff deviate significantly from Hardy-Weinberg expectations in any particular year. However, these deviations are not large, with an average inbreeding coefficient of *F_IS_* = *−*0.021 (SE = 0.004) indicating *∼* 2% inflation in heterozygosity relative to expectations under random mating. Although there is a shallow gradient of *F_IS_* with the MAF, the mean estimate is nearly identical to the average of -0.020 observed over ten midwest-US populations of *D. pulex* (Maruki et al. 2022). The central point is that temporary-pond populations of *D. pulex* have genotype frequencies that are close to the expectations under random mating, with the absolute magnitude of *F_IS_* being several-fold lower than what is commonly seen in *Drosophila* and human population samples (Johnson and Shaffer 1973; Smith et al. 1978; Prout and Barker 1993; Jin and Chakraborty 1995; Gazal et al. 2015).

**Figure 1.**
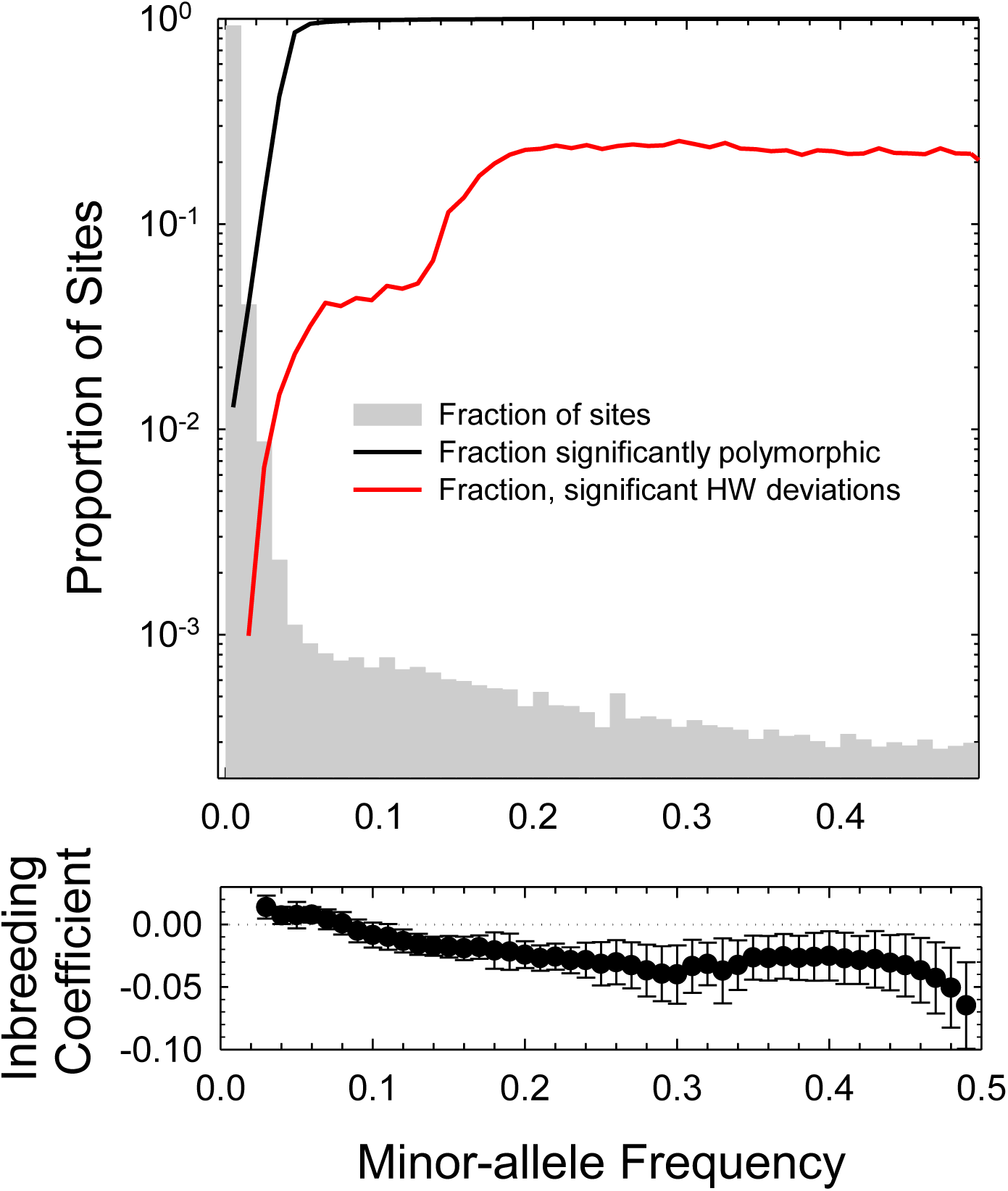
Top) Summary of the polymorphism and Hardy-Weinberg (HW) statistics, with fractions of sites with different minor-allele frequency (MAF) estimates given as the bar graph. Bottom) The average inbreeding coefficient, *Fjs,* as a function of the MAF (bars denote two standard errors); the dotted line is the expected value (0.0) under random mating.

Genome-wide nucleotide diversities, measured as the average site-specific heterozygosity under random mating (Tajima 1983), depend on the functional context, as generally seen in other organisms (Table 1). The average nucleotide heterozygosity across the entire genome is 0.0083 (SE = 0.0002), similar to that from the *D. pulex* metapopulation analysis, 0.0078 (0.0004) (Maruki et al. 2022). Silent sites and sites within a restricted region of coding introns (known to be under relaxed selective constraints; Lynch et al. 2017) have the highest levels of variation: 0.0165 and 0.0134, respectively. Replacement sites within protein-coding sequence have by far the lowest diversity (0.0037), whereas variation within UTR exons and the deeper intergenic regions (*>* 600 bp from start and stop codons) falls in the narrow range of 0.0069 to 0.0099. These sorts of relative scalings are also consistent with prior observations (Lynch et al. 2017; Maruki et al. 2022). Temporal estimates of *F_ST_*, a measure of divergence of allele frequencies on a (0,1) scale, are all very close to but significantly greater than 0.0 (Table 1). They are also *∼* 50*×* smaller than spatial *F_ST_* among *D. pulex* populations (Maruki et al. 2022), indicating that the latter estimates are not a simple consequence of variance among repeated samples.

**Table 1.** Mean within-population nucleotide diversities per site (*π*). Silent-site estimates are based on four-fold redundant sites in codons, whereas replacement sites are zero-fold redundant sites. Coding-region intron analyses span positions 8 to 34 from both ends; and intergenic analyses exclude the first and last 600 bp between the start and stop codons of adjacent genes. The 10-population (metapopulation) results are from Maruki et al. (2022). *F_ST_*estimates are based on sites with minor-allele frequencies *>* 0.1. SEs are in parentheses.

| Genomic region | Nucleotide Diversity | | $F_{ST}$ | |
| --- | --- | --- | --- | --- |
|  | PA | Metapopulation | Temporal, PA | Metapopulation |
| Silent site | 0.0165 (0.0003) | 0.0177 (0.0006) | 0.0034 (0.0001) | 0.2627 (0.0004) |
| Replacement site | 0.0037 (0.0001) | 0.0036 (0.0000) | 0.0063 (0.0001) | 0.2738 (0.0005) |
| Coding intron | 0.0134 (0.0002) | 0.0138 (0.0006) | 0.0050 (0.0001) | 0.2782 (0.0004) |
| Intergenic, Head-to-head | 0.0077 (0.0002) | 0.0080 (0.0003) | 0.0074 (0.0001) | 0.2711 (0.0003) |
| Intergenic, Head-to-tail | 0.0082 (0.0001) | 0.0082 (0.0001) | 0.0077 (0.0001) | 0.2697 (0.0002) |
| Intergenic, Tail-to-tail | 0.0073 (0.0002) | 0.0076 (0.0003) | 0.0093 (0.0001) | 0.2608 (0.0004) |
| 5' UTR exon | 0.0082 (0.0002) | 0.0085 (0.0003) | 0.0040 (0.0001) | 0.2819 (0.0005) |
| 3' UTR exon | 0.0069 (0.0001) | 0.0071 (0.0001) | 0.0045 (0.0001) | 0.2912 (0.0006) |
| 5' UTR intron | 0.0083 (0.0002) | 0.0087 (0.0003) | 0.0042 (0.0001) | 0.2840 (0.0005) |
| 3' UTR intron | 0.0099 (0.0002) | 0.0103 (0.0003) | 0.0069 (0.0004) | 0.2888 (0.0021) |

The scaling of linkage disequilibrium (LD) with physical distance between sites was determined as the squared correlation between allelic states (*r*^2^), averaged over all pairs of sites within samples and then across the nine annual samples, applying the maximum-likelihood procedure of Maruki and Lynch (2014) to sites in Hardy-Weinberg equilibrium. These estimates were acquired for different windows of allele frequency, owing to the dependence of *r*^2^ on the latter (VanLiere and Rosenberg 2008). The results are concordant with those presented for multiple populations in Lynch et al. (2022), with *r*^2^ declining in a power-law fashion up to a distance of *∼* 1 kb, and thereafter declining progressively more rapidly with distance (Figure 2).

**Figure 2.**
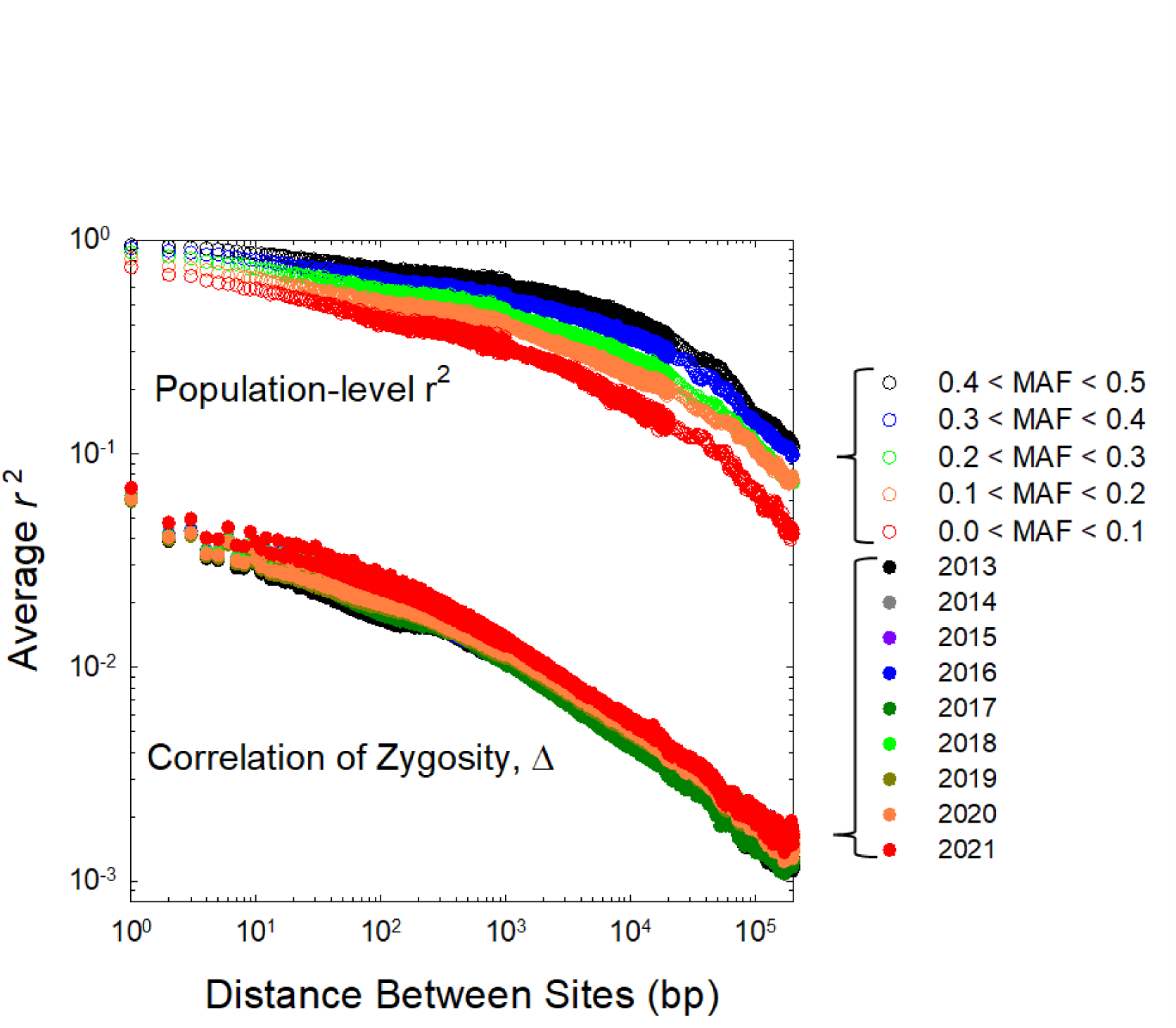
Scaling of the pattern of linkage disequilibrium (LD) with physical distance between sites, *r^2^* is the population-level LD, with averages over all sites and years given for various windows of minor-allele frequencies. Δ is the correlation of zygosity, an individual-based measure of LD based on the joint distribution of heterozygosity at pairs of sites, here given as averages over ten individuals from each of the nine sampling years.

We also determined the scaling of the correlation of zygosity (Δ; Lynch 2008), an individual based measure of LD based on the spatial distribution of heterozygous sites, for each of the ten individuals with the greatest average genome sequencing coverage in each year, using the program mlRho (Haubold et al. 2010). The average profiles for each year are highly similar to each other, and nearly parallel to the *r*^2^ profile for low-frequency MAFs (Figure 2). Given that the vast majority of polymorphisms involve low-frequency MAFs, this behavior is expected, as Δ scales approximately as *πr*^2^ (Lynch et al. 2014). These overall results indicate that evolution at individual nucleotide sites in the study species is influenced by events occurring at linked sites up to distances of *∼* 10^5^ bp, in accordance with the length distribution of “islands of strong selection” in this population (Lynch et al. 2023).

### Site-frequency spectra (SFSs)

Inferences about selection and/or demographic history associated with different categories of genomic DNA can be gleaned from genome-wide site-frequency spectra, which summarize the distribution of allele frequencies across the genome. As we do not yet have confident information on the ancestral states of alleles in this species, we resort to the folded SFS, focusing on the frequency classes for the minor alleles (i.e., with frequencies *<* 0.5). Given that between-year changes in allele frequencies are typically quite small and fluctuating in direction (Lynch et al. 2023), we pooled the total set of individuals over the course of this study to obtain average allele frequencies down to levels of *∼* 0.001 at each polymorphic nucleotide site.

Previously, from a single-year survey from a nearby population, we used the observed form of the SFS (with binned frequency classes of width 0.01) to infer that silent sites and intron sites behave in a nearly neutral manner (Lynch et al. 2017). However, with this larger data set, it is clear that the form of the SFS for even these relatively innocuous site types is inconsistent with expectations for neutral sites under mutation-drift equilibrium, which predicts an inverse scaling with *p*(1 *− p*), or 1*/p* for *p «* 1, where *p* is the minor-allele frequency (MAF) (Wright 1938; Messer 2009). Most notably, the decline in abundance in the low-frequency classes is much steeper than expected under the assumption of neutrality and constant population size, exhibiting *∼* 1*/p*^2^ to 1*/p*^2.5^ scaling up to *p ’.::* 0.01 (Figure 3A). On the other hand, beyond this point, the slope of the SFS becomes progressively weaker, converging back to the approximate pattern expected under neutrality.

**Figure 3.**
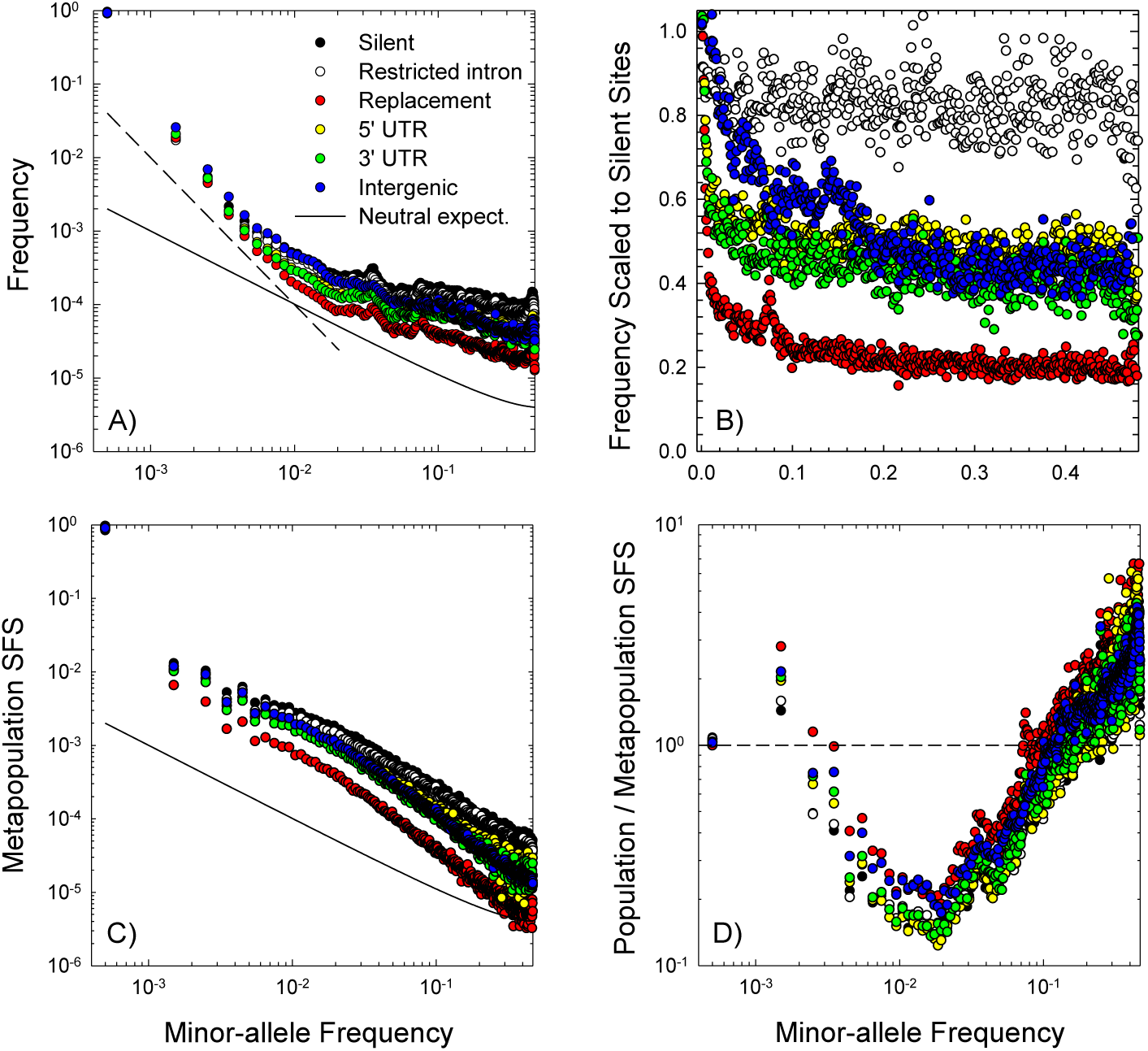
The pooled site-frequency spectra (SFS) for five classes of genomic sites. Results for silent and replacements sites are based, respectively, on for four-fold and zero-fold redundant sites. A) Results from the pooling of the nine annual PA samples; the monomorphie sites are included in the first bin. The sealing under the neutral expectation is given by the solid line, oc 1/[p(l — p)] where p is the allele frequency, with the elevation being arbitrarily set for visualization purposes. The dashed line is the expected pattern under a scaling of 1/p^2^. B) The SFS for site categories sealed by that for silent sites. C) The SFS for the ten-population pooled metapopulation sample from Maruki et al. (2022), with neutral scaling shown in panel A included for comparison. D) Ratios of the within-population (PA) SFSs to those for the corresponding site types for the metapopulation sample (i.e, the ratio of the data in panels A and C.

Three other points are apparent from the SFS profiles for alternative functional site types. First, for very low frequencies, all SFSs converge to similar *∼* 1*/p*^2^ scaling behavior (Figure 3A), as expected for alleles with low enough frequencies to be primarily governed by linked stochastic forces (including background selection). Second, 3*^1^* and 5*^1^* UTR sites appear to experience similar and intermediate strengths of selection, while selection is substantially weaker on low-frequency variants at intergenic sites but much stronger at amino-acid replacement sites. This can be seen most easily by dividing the SFSs for these selected classes by the silent-site SFS, which shows that beyond a frequency of *∼* 0.15, all profiles become relatively flat, implying that the majority of minor alleles that rise to such levels are behaving in the same way as silent-site variants (Figure 3B). Third, the within-population SFS is substantially different from the SFS for the metapopulation obtained in Maruki et al. (2022) (Figures 3C,D). Most notably, at the within-population level, there is a deficit of low-frequency alleles 0.004 to 0.1, and a corresponding inflation in the frequencies of alleles outside of this range.

What potential factors might be contributing to the form of these fine-scaled SFSs, most notably the exceptionally rapid rate of decline in allele frequencies with increasing *p* in the lowest frequency classes, *p <* 0.01? Although such results might arise if the PA population is not in demographic equilibrium (contrary to the assumptions of Equations 1a,b), this would require an extremely strong, recent population expansion, for which our prior analyses provide no support (Lynch et al. 2020; Maruki et al. 2022). If, however, there is a high level of recurrent input of background deleterious mutations of small effect, even with a constant population size, the SFS for neutral sites is expected to have a very steep initial slope followed by a gradual return to the neutral-expectation scaling (Cvijovíc et al. 2018). This happens because most new mutations arise linked to segregating deleterious mutations at one or more sites, which eliminates neutral mutations more rapidly than expected by chance (Charlesworth 2012), with rising to high frequencies residing on relatively clean linked backgrounds. As pointed out by Cvijovíc et al. (2018), this predicted effect will not be seen unless the SFS is based on large enough numbers of individuals to prevent the signature behavior from being obscured within a single low-frequency bin, and indeed the predicted pattern has not been previously reported, including in our own work (Lynch et al. 2017; Maruki et al. 2022), which involved sample sizes of *<* 100 individuals per population.

With bins of width 0.001, the SFSs illustrated in Figure 3A are qualitatively consistent with a strong influence from background selection. The theory predicts an inverse scaling of the SFS with *p* for *p* smaller than *p*_1_ = 1*/*(*N_e_^√^U_d_s_d_*), where *N_e_*is the population size, *U_d_* is the haploid rate of deleterious mutation within regions effectively fully linked to the neutral markers, and *s_d_* is the average deleterious effect of linked mutations. Prior work with *D. pulex* suggests *U_d_s_d_:::* 0.002 to 0.028 at the whole-genome level (Deng and Lynch 1997; Lynch et al. 1998; Schaack et al. 2013; Latta et al. 2013). It is unclear how much smaller *U_d_s_d_* will be within the span of strong linkage blocks, but assuming *U_d_s_d_* is as large as 10*^−^*^4^ (a 100-kb linkage block being *∼* 0.05% of the genome), the upper bound on *p* required for inverse scaling would still be *p*_1_ ::: 100*/N_e_*. Given that the estimated effective size of the PA population is of order 10^6^ (Lynch et al. 2020), the lowest frequency class that we observed (::: 0.001) is likely still beyond the domain of effective neutrality (*p <* 10*^−^*^4^). More notable is the prediction that for frequencies within the range of *p*_1_ up to a second cutoff defined by *p*_2_ = *e^U^^d/sd^ /*(*N_e_s_d_*), the SFS is expected to scale inversely with *p*^2^, then gradually returning to *p^−^*^1^ scaling, in qualitative accord with the observed data (Figure 3A). As the theoretical expectations derived in Cvijovíc et al. (2018) were based on the assumption of mutations with constant effects, and the actual distribution of fitness effects likely spans orders of magnitude, the scaling bounds noted above are only approximations.

A second factor that may contribute to the steep decline the SFS for low-frequency MAFs is fluctuating selection, i.e., temporal variation in selection coefficients, which can distort the form of the SFS towards alleles with extreme frequencies. We know from prior work (Lynch et al. 2023) that nucleotide sites throughout the genome are under weak selection with temporal variation in magnitude between years. The average net selection coefficient (from both direct and linked effects) for minor alleles is in the range of -0.002 to -0.007 (with silent sites and intron sites at lower end of the range), whereas the average temporal variance (among years) of selection coefficients ::: 0.16. Considerable attention has been given to the influence of fluctuating selection on the form of the SFS (e.g., Wright 1948; Karlin and Levikson 1974; Takahata and Kimura 1979; Dean 2018), the most complete derivation having been developed by Taylor (2013). The critical determinants of the SFS are the composite parameters 4*Nu*, 4*Nσ*^2^, and *s/σ*^2^, where *s* and *σ*^2^ are the temporal mean and variance of *s*.

One concern with the underlying theory is that *N* is assumed to be the effective population size determined by factors extrinsic from the fluctuating-selection process. Therefore, to more thoroughly evaluate the influence of fluctuating selection on the SFS, we performed computer simulations of a single biallelic nucleotide site experiencing temporally fluctuating selection (using a Gaussian distribution of selection coefficients, with each of the two alleles assumed to have the same *σ*^2^, while allowing for different allele-specific average *s*, and *s* being defined as the difference between the two) and recurrent reversible mutation in an ideal Wright-Fisher population of size *N* = 10^6^, and a per-site mutation rate of 5.7 *×* 10*^−^*^9^ (Keith et al. 2016). Along with the selected site, the SFS of a completely linked neutral site was monitored over a long enough time scale to generate the steady-state distribution of frequencies of both selected and neutral sites.

Fluctuating selection has two potential effects on standing levels of diversity. First, from the standpoint of the selected site, if *Ns <* 1, the SFS behaves as though the sites are essentially neutral, even for *Nσ*^2^ *»* 0 (Figure 4). At the opposite extreme, if *Ns >* 1 and *σ*^2^*/s «* 100, the directional power of selection dominates the selection process, and few deleterious alleles ever arise to high frequency. If, however, *Ns >* 1 but *σ*^2^*/s >* 100, the fluctuating-selection forces dominate the directional power of selection, and the SFS again has a form close to the neutral expectation, with a long plateau to the right. As the latter condition appears to be frequently met in the PA population (Lynch et al. 2023), this means that the strong plateaus in the SFSs for the groups of functionally significant sites need not be simply due to heterogeneity in selection coefficients among sites (i.e., to the subset of alleles surviving to high frequencies being effectively neutral).

**Figure 4.**
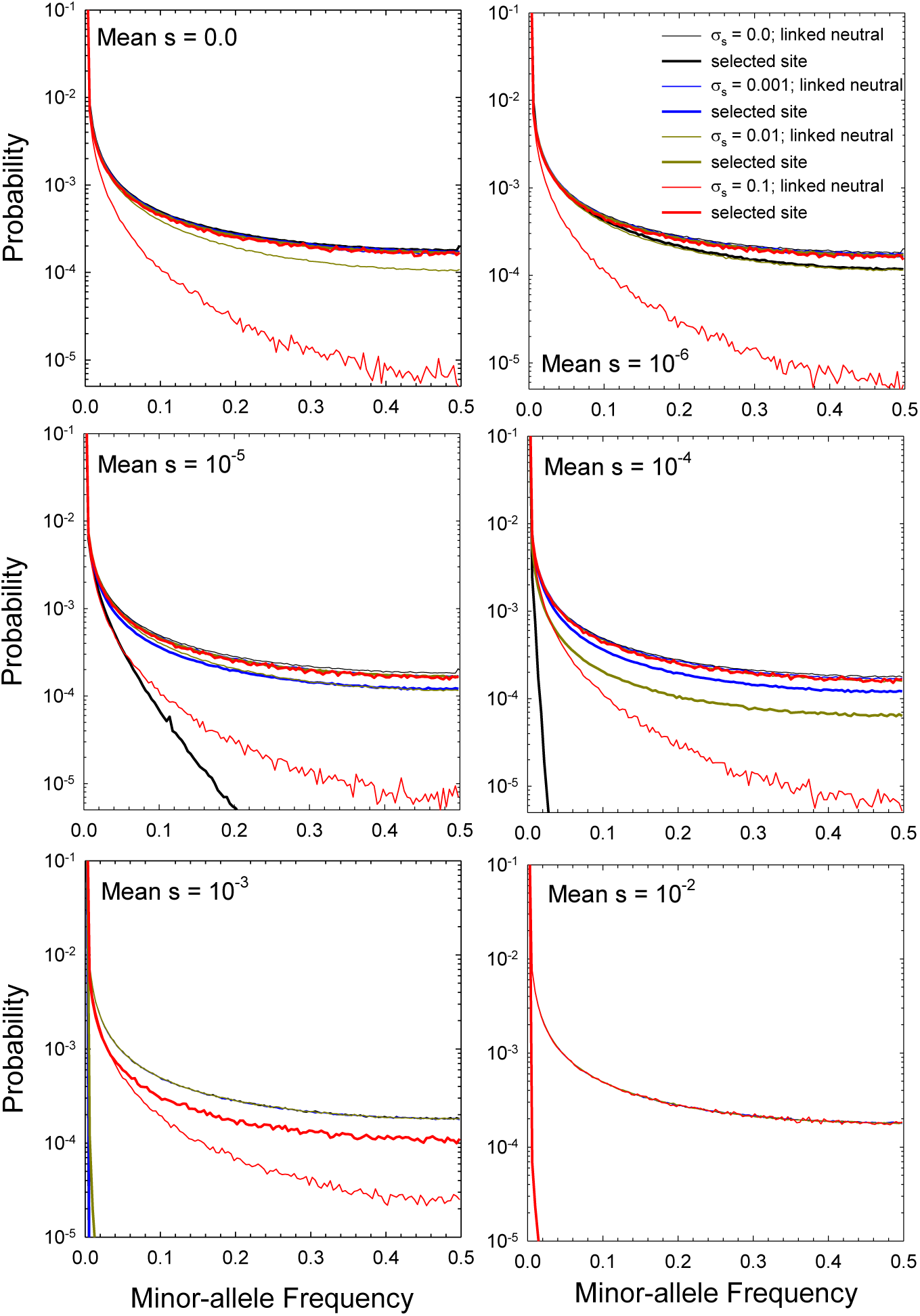
Forms of the site-frequency spectra (SFSs) for selected sites and linked neutral sites, for various levels of J and σ_s_, derived by simulations in a Wright-Fisher format as described in the text. Each folded SFS is based on 250 frequency bins of width 0.004, with counts derived from simulations generally over 10^tì^jV generations, where Λ^ľ^= 10^G^ is the population size. With very strong mean selection (s), the results for selected sites are generally not visible, as the probability of attaining appreciable frequencies is very close to 0.0.

Second, as anticipated by Taylor (2013), fluctuating selection can alter the form of gene genealogies in ways that can modify the standing level of variation at linked silent sites. In effect, fluctuating selection has two opposing effects. Haplotypes with selected-site frequencies near 0/1 have an elevated probability of loss/fixation, whereas those at intermediate frequencies experience a form of balancing selection. The overall effect is that the depth of haplotype gene genealogies will be reduced, thereby diminishing the expected variation at silent sites, while retaining a configuration reflecting an elevated probability of maintenance of a balanced polymorphism at the selected site. The simulation results show that provided *s ≤* 0.01, for *Nσ*^2^ *>* 100, the height of the SFS for linked neutral sites is substantially depressed (up to tenfold for high-frequency alleles). This leads to situations in which the probabilities of intermediate frequencies for the selected site are elevated relative to the case at linked silent sites, thereby resulting in ratios of amino-acid replacement-site to silent-site diversities, *π_N_ /π_S_*, greater than the neutral expectation of 1. When *s* is very large, this effect is eliminated because the selected locus is nearly always fixed for the beneficial allele, which with 4*Nu «* 1 renders the form of the gene genealogy (the coalescent) close to the neutral expectation.

### Polymorphism in protein-coding genes

To obtain insight into the degree to which selection influences standing variation at protein-coding loci, at each of the nine sampling dates, gene-specific estimates were obtained for within-species variation at 4-fold and 0-fold redundant sites (*π_S_* and *π_N_*) as well as for intronic and intergenic regions, confining the analyses to the 13,478 genes for which there was adequate coverage data for at least four sampling years (Supplemental File S1). By averaging over all genes and years, several general statements can be made with respect to patterns of variation over the lengths of gene bodies (Figure 5). First, intergenic DNA harbors average levels of polymorphism that are nearly independent of position with respect to the start and stop codons of genes, lower than levels observed for silent sites, but higher than those for amino-acid replacement sites. Second, although replacement-site variation is also relatively constant across gene bodies, silent-site variation increases with distance from start and stop codons, reaching a constant average level only beyond 100 bp. Third, variation at sites within introns declines with increasing distance from intron-exon junctions, eventually asymptoting at essentially the same level as observed for intergenic DNA, implying (as noted above based on SFS observations) the operation of significant selection across the full lengths of introns.

**Figure 5.**
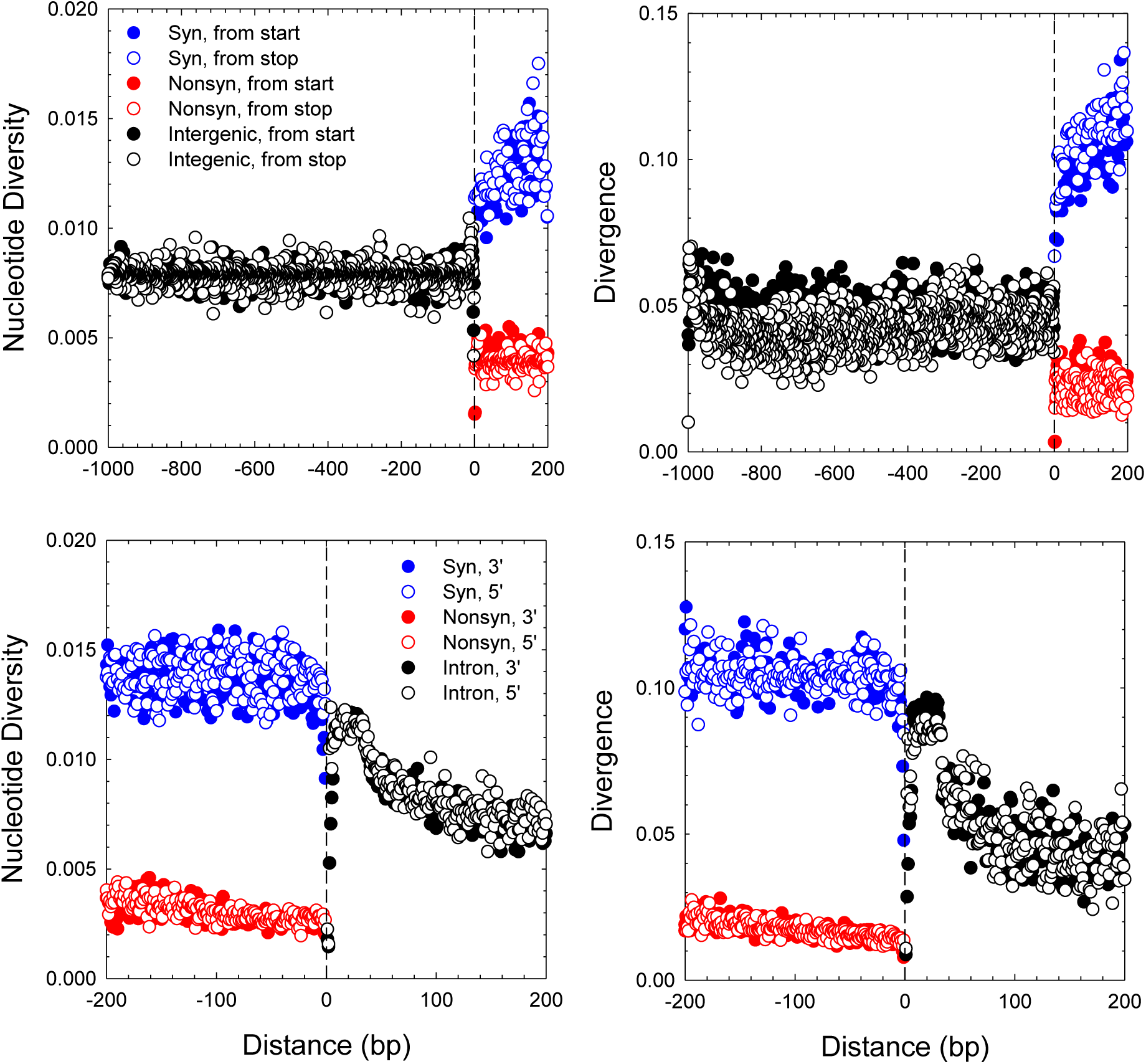
Measures of average levels of nucleotide diversity (left panels) and divergence (from *D. obtusa* orthologs (right panels). For the upper panels, site positions are given in absolute locations downstream of translation start codons and upstream of stop codons, with the estimates for each gene extending just to the half distance to the nearest intron. For the lower panels, site positions are given in absolute distances from each end of introns, with estimates for exons extending to the midpoint of the distance to the next intron, and estimates for introns extending to intron midpoints.

Additional downstream gene-specific analyses were subdivided into genes 11,853 and 1,625 with and without clear orthologs in the outgroup *D. obtusa,* respectively. With multiple independent (annual) estimates of each gene-specific parameter, overall averages and standard errors of the two diversity parameters were obtained from the individual sample estimates, and the *π_N_ /π_S_* index was estimated as the ratio of *π_N_*to *π_S_*, with its standard error obtained by applying the sampling variances of the components to the expression for the variance of a ratio (Lynch and Walsh 1998).

After removing 35 outliers with ratios *>* 10.0 (likely due to spuriously low estimates o *π_S_*), the average *π_N_ /π_S_* over all genes with *D. obtusa* orthologs is 0.257 (SE = 0.006, median = 0.131, mode = 0.033; Figure 6A), somewhat higher than the average over the 10-population study (Table 2). The 1,625 genes without *D. obtusa* orthologs have a highly elevated mean *π_N_ /π_S_* of 1.031 (SE = 0.033, median = 0.582, mode = 0.280), not significantly different from the neutral expectation of 1.0 (after removing 71 outliers with ratios *>* 10.0). The subset of putative PA genes without *D. obtusa* orthologs are relatively short in length, averaging 408 (SE = 7) codons, which is *∼* 68% of the average of 597 (5) for genes with *D. obtusa* orthologs. Although these observations suggest that the genes without *D. obtusa* orthologs are evolving in a neutral fashion, the mean *π_N_ /π_S_ :::* 0.324 for them in the metapopulation study, suggesting that many of them are likely to be functionally significant in some populations and/or years.

**Figure 6.**
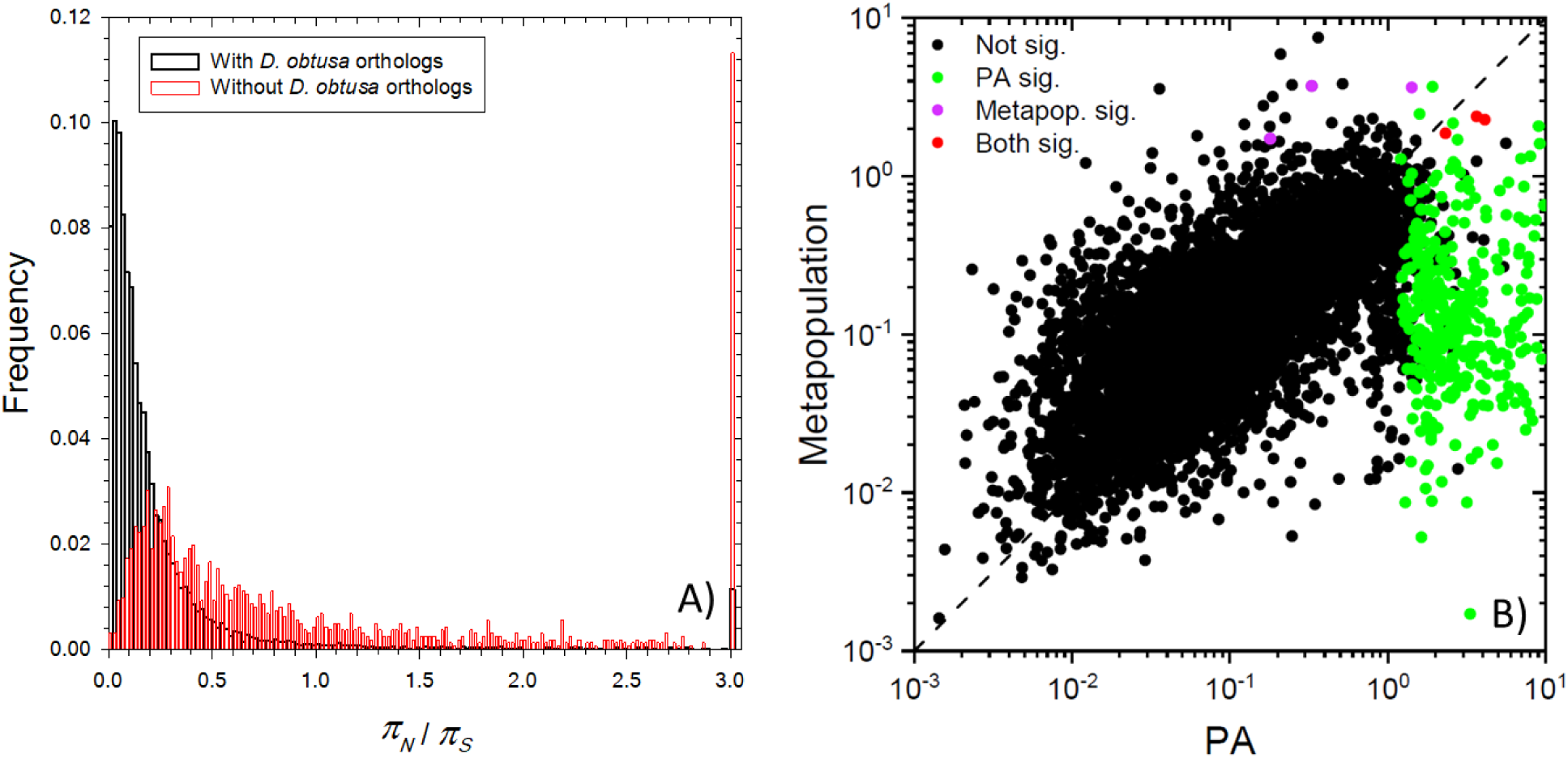
Frequency distributions of the polymorphism ratio, kw/tγs. A) πv/πs for classes of genes with and without orthologs in the outgroup species D. øϋΐωα, B) Joint distributions of 7ΓĄrƒτr_5_ for the PA population and the average of ten Zλ jow/ea? populations (from Maruki et ah 2022). The diagonal dashed line denotes equality. Colored points denote estimates that arc significantly > 1.0.

**Table 2.** Comparison of sequence diversity and selection indicators for 11,853 protein-coding genes with and 1,625 genes without orthologous data in *D. obtusa.* The metapopulation results are from Maruki et al. (2022). SEs are in parentheses.

|  | PA Only | Metapopulation |
| --- | --- | --- |
| With *D. obtusa* orthologs: |  |  |
| $\pi_N$ | 0.0026 (0.0000) | 0.0024 (0.0000) |
| $\pi_S$ | 0.0175 (0.0001) | 0.0158 (0.0001) |
| $\pi_N/\pi_S$ | 0.2565 (0.0056) | 0.1908 (0.0026) |
| $d_N$ | 0.0219 (0.0004) | 0.0165 (0.0001) |
| $d_S$ | 0.1367 (0.0005) | 0.1225 (0.0004) |
| $d_N/d_S$ | 0.1481 (0.0017) | 0.1397 (0.0012) |
| Without *D. obtusa* orthologs: |  |  |
| $\pi_N$ | 0.0060 (0.0001) | 0.0040 (0.0001) |
| $\pi_S$ | 0.0133 (0.0003) | 0.0184 (0.0003) |
| $\pi_N/\pi_S$ | 1.0313 (0.0331) | 0.3243 (0.0084) |

Out of the total pool of 13,478 genes analyzed, 1,008 have *π_N_ /π_S_ >* 1.0, a potential signature of balancing selection at the amino-acid level. The subset of such genes with *π_N_ /π_S_* significantly *>* 1.0 were identified as cases in which the estimate of *π_N_* exceeded that for *π_S_* by more than two standard errors. Using a one-tailed Z-test followed by Bonferroni correction for multiple comparisons, 583 such genes are significant at the 0.05 level, 335 of which have no orthologs in *D. obtusa.* Strikingly, 381 (65%) of these high-*π_N_ /π_S_* outlier genes are located on chromosome 12 (Supplemental Figure S1), almost all on one arm. In the metapopulation study, only 28 genes exhibited average *π_N_ /π_S_* significantly greater than 1.0 (Maruki et al. 2022), only six of which were also highlighted in this study, and only two of which are on chromosome 12. This suggests that a moderate amount of balancing selection may be occurring in individual populations, with different loci being involved in different ecological settings. Chromosome 12 in the study population might be one such example, although its *F_IS_* = *−*0.0157 (0.0004) is less extreme than the genome average, *−*0.0208 (0.0001).

Notably, the pool of 583 high-*π_N_ /π_S_* genes has an average *π_S_* = 0.0024 (0.0001), substantially below the global average across the genome, 0.0169 (0.0001). In contrast, average *π_N_*, 0.0063 (0.0002), is far greater than the genome-wide mean of 0.0026 (Table 2), so the elevated *π_N_ /π_S_* is due to both of these features. These observations are consistent with the predictions of fluctuating-selection theory noted above. GO enrichment analysis revealed that the overall pool of PA genes with *π_N_ /π_S_* significantly *>* 1.0 is enriched in several categories involving DNA/RNA processing, neuro-transmission, development, and lysosome activity (Supplemental File S1).

The PA gene estimates for *π_N_ /π_S_* are moderately correlated with those obtained in the metapopulation study of Maruki et al. (2022) (*r*^2^ = 0.468 and slope = 0.583 (SE = 0.007), on a log-log scale) (Figure 6B). For the 9,210 genes evaluated in both studies, 331 were significantly *>* 1.0 at the 0.01 significance level in the PA population, whereas 6 were so in the metapopulation study, and only 3 were jointly significant. These results highlight the substantial sampling variance associated with these kinds of diversity estimates even with haplotype sample sizes exceeding 1000. Excluding outliers with *π_N_ /π_S_ >* 10, the average coefficient of sampling variance (SE/mean) of *π_N_ /π_S_* is still 0.131 (SD = 0.102) in the current study.

### Between-species divergence

To evaluate levels of interspecific divergence, we compared the 11,853 one-to-one orthologs with *D. obtusa,* which has an average 0.1367 (0.0005) divergence at silent sites (*d_S_*) with respect to the PA population. Average patterns of divergence over gene bodies are very similar to those noted above for within-species variation (Figure 5). Noting that the level of divergence between two species for neutral sites ::: 2*µt*, where *µ* is the base-substitution rate per nucleotide site per generation, and *t* is the number of generations since reproductive isolation, then under the assumption that silent-site variation is effectively neutral, with expected value 4*N_e_µ*, the ratio of divergence to polymorphism for such sites has an expected value of *t/*(2*N_e_*) generations. For silent sites 3*^1^* and 5*^1^* relative to intron-exon junctions, the average ratios are 7.65 (0.05) and 7.54 (0.04), respectively. Thus, across the genome, the average degree of isolation between *D. pulex* and *D. obtusa* is *∼* 15*N_e_*generations.

The average *d_N_ /d_S_* for this species pair is 0.1481 (0.0017), substantially lower than average *π_N_ /π_S_* = 0.2565 for this particular subset of genes (Table 2). Only 95 genes have *d_N_ /d_S_* exceeding the neutral expectation of 1.0, with just 54 of these exceeding this benchmark at the 5% confidence level after correction for multiple comparisons, and 12 of the latter having no apparent orthologs outside of *Daphnia* (Supplemental File S1). Again, we found that most (44 of 54) of theses outlier genes are located on chromosome 12. Only one outlier, LOC124207880, overlaps with the pool of *d_N_ /d_S_ >* 1 genes identified as significant in the prior metapopulation study, and it is unannotated and hence with unknown function.

As *d_N_ /d_S_ >* 1 is a highly conservative measure of positive selection, we evaluated the neutrality index (NI), the ratio of *π_N_ /π_S_* to *d_N_ /d_S_*, to search for genes with excess aminoacid divergence in the PA population relative to expectations based on polymorphisms. NI is expected to take on values *<* 1.0 when there is substantial positive selection for protein-sequence divergence at the between-species level, and values *>* 1.0 when selection is primarily purifying in nature. About 2.7% of annotated genes have NI in excess of 25.0 in the PA population, but many of these are extremely short (and possibly false annotations), which may cause spurious behavior of ratio estimates. Ignoring genes with NI *>* 25.0, the mean NI is 2.008 (0.0027) significantly above the neutral expectation of 1.0, indicating that within-population purifying selection is the predominant form of selection operating on amino-acid sequences. However, of the 11,693 genes in this analysis, 39% have NI *<* 1.0, 2,038 (*∼* 17%) of which have NI significantly *<* 1.0 at the 5% probability level after correction for multiple comparisons (Supplemental File S1), suggesting widespread positive selection (in one or both species).

There is a strong correlation between NI estimates obtained in this study and in the prior metapopulation analysis, which together comprise *>* 3, 000 sequenced haplotypes (Figure 7). To evaluate the classes of genes harboring the strongest signature of positive selection (for divergence), we focus on 499 candidate genes derived from three sets of outliers: 318 genes exhibiting NI significantly *<* 1.0 in both studies; 42 genes exhibiting significantly low NI only in the metapopulation study, but with the mean from both studies remaining significant; and 139 genes exhibiting significantly low NI in the PA study, with the mean from both studies remaining significant (Supplemental File S2). In our prior work, we identified chromosomal islands with evidence of strong directional selection over the time course of this study (Lynch et al. 2023), but we find no evidence that the 499 candidate genes here are enriched in such regions, implying that the outcome of long-term selection underlying interspecies divergence confers no information on short-term selection processes. However, the pool of 499 outliers is enriched on chromosome arms (as opposed to centromeric regions). Although centromeric regions constitute 42.2% of the entire genome, only 16.8% (84) of the outlier genes are located in such regions (Supplemental Figure S2).

**Figure 7.**
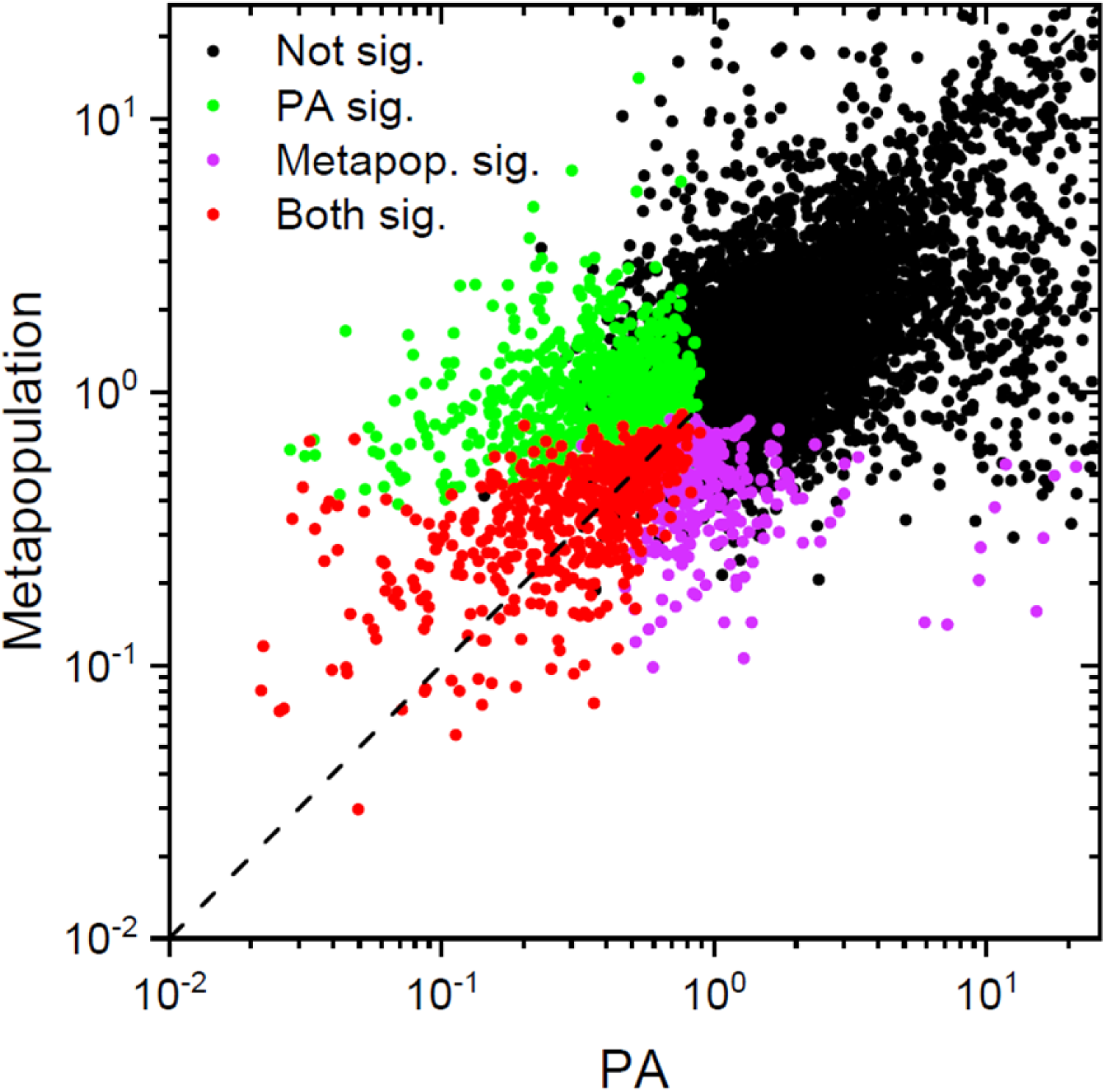
Joint distribution of neutrality-index estimates from nine years of PA data and from a mctapopulation study; *r^2^ —* 0.267; slope= 0.373 (SE = 0.007); Λ^τ^ = 8078). Colored points denote estimates that arc significantly < 1.0.

Considering the set of 100 genes with the lowest NI estimates (all but nine of which were jointly significant in both studies), NI ranged from 0.040 to 0.219 (average SE = 0.055), implying a level of divergence between 25 and 5*×* the expectation based on polymorphism data (Supplemental File S2). Based on homology to known genes across the Tree of Life, this pool of extreme outliers is enriched for genes associated with ribosomal proteins (5 cytosolic and 7 mitochondrial), nuclear-encoded mitochondrial proteins (12, including those involved in bioenergetics), and neurobiology (14). Three of the four most extreme low-NI outliers are associated with mitochondrial ribosomes. Of the 20 most extreme genes, four are associated with ubiquitinylation processes (which typically mark proteins for degradation), and two with lifespan determination in metazoans (N-alpha-acetyltransferase; and the gene with the lowest NI, a subunit of the mTOR complex). However, 11 of the top 100 have no known orthologs outside of Daphnia.

As a second approach to revealing the biological features experiencing the greatest intensity of positive selection, we considered the degree to which genes in the aforementioned list of 499 were enriched in various Gene Ontology (GO) categories (Supplemental File S2). Ribosomal protein-coding genes were again identified as the most enriched category of rapidly evolving genes (with the mitochondrial proteins being particularly enriched). Also highly enriched were genes associated with other aspects of translation, extracellular matrix, ion channels and synaptic vesicles, and electron transport in the mitochondrion (10*^−^*^5^ *< P <* 10*^−^*^4^). Moderately enriched categories (10*^−^*^4^ *< P <* 10*^−^*^2^) include RNA polymerase III subunits (involved in transcription of ribosomal 5S rRNA, tRNA and other small RNAs), neuronal systems (including opsins used in light perception, and ion transport at neuronal junctions), other aspects of mitochondrial biology, and other aspects of the nervous system.

In a third approach, we identified the GO categories with the lowest mean NI (over the total pool non-outlier plus outlier genes). These include: 1) biological processes associated with lipid biosynthesis, RNA processing, and developmental signaling; 2) cellular components associated with organelle membranes, neuronal development, maintenance of DNA integrity, microtubules, and motors; and 3) molecular functions associated with transmembrane transport, neurotransmission and neuroendocrine response, and cell-cell adhesion (Figure 8). Most of these extreme groupings still have an average NI *>* 0.7, and in some cases *>* 1.0, indicating that even the GO groups with the highest levels of positive selection still contain substantial fractions of highly conserved genes. However, the consistent message from these three analyses is that the genes under the strongest level of positive selection in the *D. pulex* lineage tend to be associated with mitochondrial and cytosolic ribosomes, other mitochondrial functions, other aspects of translation, and the nervous system.

**Figure 8.**
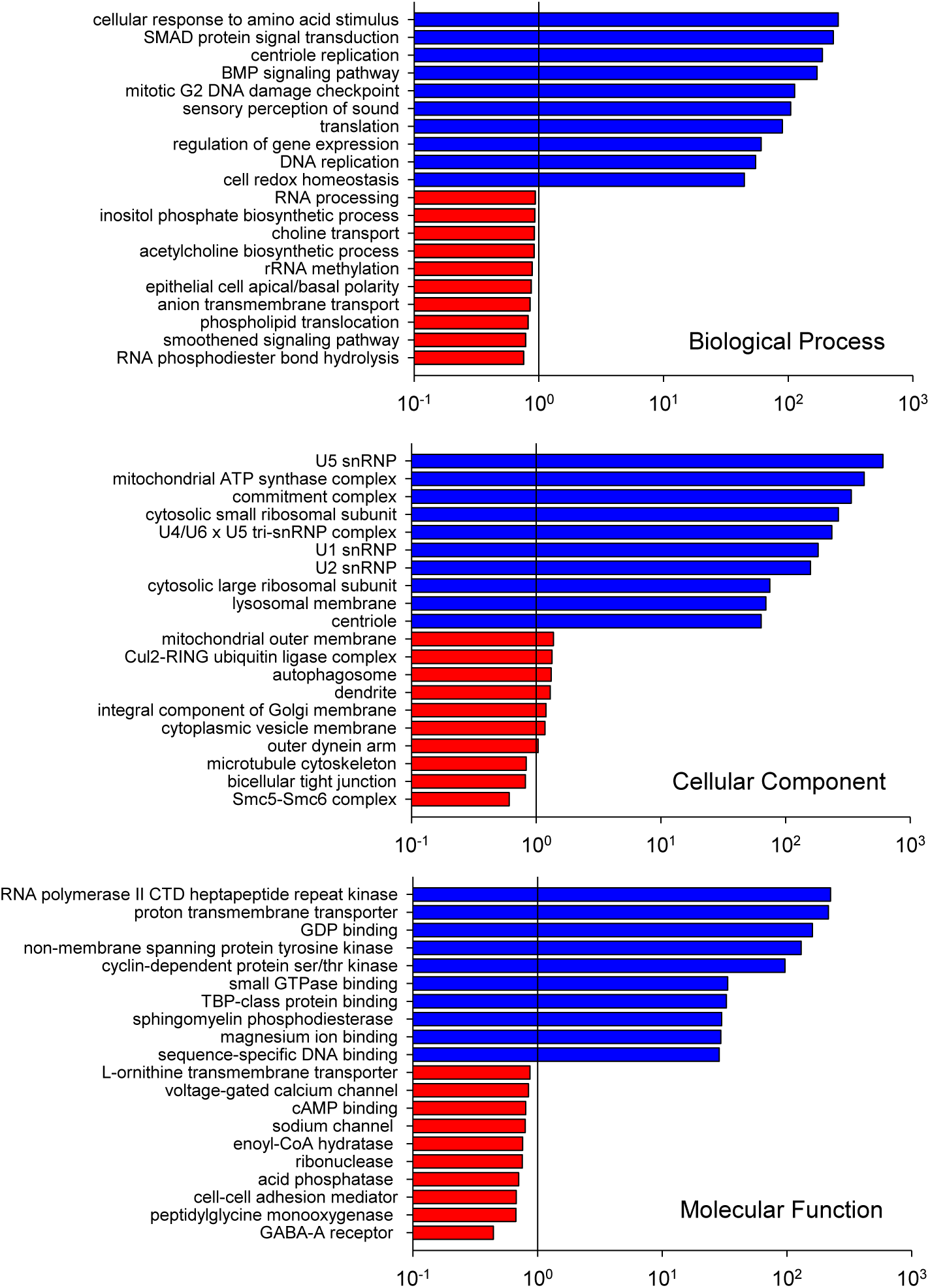
Gene-ontology categories with the highest and lowest average NI indices over all component genes, denoting the strongest indications of purifying (blue) vs. positive (red) selec­tion. Results arc given for the ten outliers at the extremes of the distributions for the primary categories of biological processes, cellular components, and molecular functions.

Finally, we consider the genes with NI significantly greater than 1 (again, based on a summary from both the PA and metapopulation studies), i.e., the subset of genes experiencing the strongest levels of purifying selection. Of the 632 genes that met this criterion, 44 have no identifiable orthologs outside of *Daphnia* (eukaryote or prokaryote). (Supplemental File S3). These genes have an average *π_N_ /π_S_* = 0.475 (SE = 0.060), nearly double that for genes with non-*Daphnia* orthologs (above), which in part accounts for their high NI. However, as only 10% of such genes have *π_N_ /π_S_ >* 1 (the neutral expectation), it appears that most of these unknown genes are under purifying selection and hence of biological significance to *Daphnia.* The average *d_N_ /d_S_* for this subset, 0.196 (SE = 0.015), is *∼* 30% greater than that for the full pool of *D. obtusa* orthologs, so the elevated NI is primarily a consequence of excess polymorphism for amino-acid altering mutations.

For the high-NI outlier genes with inferred orthologs outside of *Daphnia,* a substantially different picture emerges than found for the low-NI outliers (Supplemental File S3). This pool of relatively slowly diverging genes is dominated by those with functions in growth and metabolism, regulation and post-translational modification, cytoskeletal functions, and extracellular matrix. Contrary to the situation with the low-NI outliers, genes associated with ribosomal proteins are rare, and the few that are associated with mitochondria have functions associated with membrane transport and metabolic functions (not associated with the electron transport chain or ribosomes). The primary GO enrichment categories that are significant at the *P <* 0.01 level for high NI are: 1) biological processes associated with heterochromatin assembly, transmembrane transport, mitochondrial organization, mitotic DNA replication checkpoint, microtubules, protein glycosylation, and ubiquitin-dependent ERAD pathway; 2) cellular components associated with kinetochores, Golgi, mitochondrial outer membrane, and cilia; 3) molecular functions associated with flavin adenine dinucleotide (FAD) binding, protein serine/threonine phosphatase activity, RNA helicase activity, microtubule binding and motor activity, and DNA polymerase.

GO categories with the highest mean NI (lowest relative divergence rates) have: 1) biological processes associated with growth regulation, genome replication, gene-expression regulation, translation, and redox homeostasis; 2) cellular components associated with spliceosomes, ribosomes, lysosome membranes, ATP synthesis, and centrioles; and 3) molecular functions associated with phosphorylation, membrane transport, conversion of lipids, and small-molecule binding (Figure 8). The overall interpretation here is that the most slowly diverging proteins in *D. pulex* are generally associated with chromosomal maintenance and replication, transcript processing, mitochondrial membrane proteins, energy production (redox reactions and ATP synthesis), tubulin-related cytoskeletal functions, degradation of misfolded proteins, and postranslational modification of proteins.

## Discussion

Based on an exceptionally large set of genome-wide population-genetic surveys, the results of this study highlight some of the challenges and benefits of expansive surveys of population-genomic patterns of variation. For example, although it has become common to use information on patterns of standing variation, including the SFS, to infer past demographic history (reviewed in Johri et al. 2022), our empirical results, combined with recent theory, highlight the substantial limitations of this approach. The basic features of genetics and ecology ensure that essentially all populations will persistently experience both background selection and fluctuating selection, although *D. pulex* is the only species for which the relevant quantitative information is available on both phenomena. Studies of both factors require substantial investment in long-term surveys: temporal surveys of natural populations to obtain estimates of the stochastic aspects of selection (Lynch and Ho 2020; Lynch et al. 2023); and laboratory mutation-accumulation experiments to obtain mutational parameters relevant to understanding issues related to background selection (Lynch et al. 1999; Baer et al. 2007; Katju and Bergthorsson 2019).

Moreover, although substantial theory has been devoted to both background selection and fluctuating selection, the consequences of their joint operation remains unexplored. This too is a significant issue, as the two processes are unlikely to be independent. The net effects of selection operating on individual nucleotide sites is a function of direct site-specific functional effects as well as an indirect consequence of the effects of all other tightly linked sites. Thus, the stochastic appearance of background (linked) mutations is likely to be intertwined with temporal variance in selection coefficients operating on individual nucleotide sites.

Our results raise difficult questions about the reliability of static observations of genomewide features such as SFSs to infer the genetic features of even the most well-characterized natural populations. A full characterization of the effects of background selection on the SFS for even neutral sites requires information on the rate of mutational input and distribution of effects of unconditionally deleterious mutations as well as the rate of recombination (with and without crossing over). Likewise, the effects of fluctuating selection involving conditionally beneficial/detrimental mutations brings in several more key parameters (minimally, the recurrent, reversible mutation rates for alternate alleles; their average effects; the temporal pattern of variation in selection coefficients; and the degree to which the latter two covary). Combined then, even in a demographically stable population, the joint effects of positive and fluctuating selection are functions of at least eight parameters (ignoring additional complexities such as dominance and epistasis), raising significant problems with identifiability of alternative models based on inference from a SFS. It is already known, for example, that the simplest form of fluctuating selection can yield a SFS that is essentially the same as the expectation under neutrality (Tier et al. 1982), and this is confirmed by the simulation results in Figure 4. Then, there is the additional problem that some of the most informative features of an SFS reside in the lowest frequency classes (Cvijovíc et al. 2018). Quantitative information on the abundance of these rare alleles is beyond the reach of most studies, even one as large as that herein (containing *∼* 1500 haplotypes).

These caveats aside, large-scale genomic studies of polymorphism and divergence do harbor considerable power to identify genes and gene categories under extreme forms of selection. In pursuing these issues, combining the results from a prior metapopulation survey, we took three different approaches to identifying genes experiencing exceptionally strong positive vs. purifying selection at the species-wide level: 1) identification of the most extreme gene-specific values; 2) identification of the GO categories enriched with such genes; and 3) evaluation of the overall mean estimates of the predominant direction of selection within GO categories. This complementary approach leads to the robust conclusion that the study population is experiencing particularly strong positive selection for change in nuclear-encoded proteins involved in mitochondrial ribosomes and the electron-transport chain, as well as nervous-system function, likely for very different reasons – in the former case, coevolutionary drive associated with the accumulation of mildly deleterious mutations in the rapidly mutating mitochondrial genome; and in the latter case, challenges associated with external ecological pressures.

## METHODS

### Sample preparation and sequencing

*D. pulex* individuals were collected from the same temporary pond in Portland Arch (PA), USA (40.2096, -87.3294) each year. As in Lynch et al. (2017), to maximize the likelihood that each individual would originate from a unique resting egg, we collected hatchlings in the early spring before the occurrence of subsequent reproduction. Individual isolates were clonally isolated in the laboratory for two to three generations, and from these pools, DNA was extracted from 96 isolates per population, with each prepared into a library using a Bioo or Nextera kit, followed by tagging with unique oligomer barcodes. Paired-end short (100 or 150 bp) reads were sequenced using the Illumina HighSeq 2500 platform, and adapters sequences were trimmed using Trimmomatic (version 0.36) (Bolger et al. 2014), and then mapped to the current high-quality reference assembly for a clone taken from an adjacent population (clone KAP4; NCBI accession number GCF 021134715.1) using Novoalign (version 3.02.11) with the “-r None” option. We then filtered out low-quality sites and individuals following Maruki et al. 2022: 1) removed clones with mean coverage over sites *<* 3*×*; 2) removed clones that are highly related (e.g., pairs of full sibs that might have hatched from a single resting egg); 3) removed clones with *>* 3% of asexual markers (Ye et al. 2021); 4) eliminated sites with nonbinomial deviations in read distributions identified in *≥* 4 deviant individuals using a goodness-of-fit test (Ackerman et al. 2017).

### Standing variation and inbreeding coefficients

To measure levels of standing variation, we first prepared nucleotide-read quartets (counts of A, C, G, and T) for each clonal isolate, necessary for the population-genomic analyses, using the proview command of MAPGD (version 0.4.26) (Ackerman et al. 2017). Then, allele frequencies, genotype frequencies, significance of the polymorphisms, and genotypic deviations from Hardy-Weinberg equilibrium in each sample were estimated using the package GFE (version 4.3) (Maruki and Lynch 2015). Nucleotide diversities, measured as the average site-specific heterozygosity, were obtained using the formula of Tajima (1983). *F_IS_* (mean inbreeding coefficient within populations) was also estimated using package GFE (version 4.3) (Maruki and Lynch 2015), restricting analyses to sites that were significantly polymorphic in at least one of the populations at the 5% level.

### Temporal fixation indices

Wrights (1951) fixation indices were estimated from the genotype-frequency estimates at SNP sites derived with the method of Maruki and Lynch (2015), restricting analyses to sites that were significantly polymorphic in at least one of the years at the 0.05 probability level. The framework of Weir and Cockerham (1984) was used to obtain site-specific estimates of *F_ST_*(temporal genetic differentiation among population samples), as described in Weir (1996). The method of Weir and Cockerham (1984) estimates fixation indices from accurate genotypes with no missing data. However, for data generated by high-throughput sequencing, depths of coverage vary among sites, individuals, and chromosomes within diploid individuals, so adjustments are needed to account for variability of sample sizes. We did so by estimating the effective number of sampled individuals (*n_ei_*) for which both chromosomes are sequenced at least once (Maruki and Lynch 2015) at each site within each sample. For sample *i*,

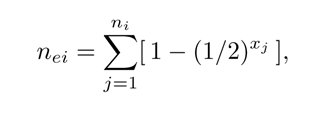

where *n_i_* and *x_j_* are the number of sampled individuals in deme *i* and depth of coverage in individual *j*, respectively. We required that *n_ei_* be at least 10 in each sample for a site to be included in the analysis. Letting *r_e_* denote the number of samples that satisfied this condition, the fixation indices were estimated by substituting *n_ei_* and *r_e_* for the numbers of sampled individuals in deme *i* and the number of demes, respectively, in the Weir and Cockerham equations.

### Patterns of linkage disequilibrium

Population-level linkage disequilibrium (*r*^2^) was estimated using the maximum-likelihood method of Maruki and Lynch (2014), which is designed for populations in HardyWeinberg equilibrium, a condition closely approximated by the study population (Maruki et al. 2022). The input for the method is the read quartet (numbers of inferred A, C, G, and T nucleotides) at each filtered site in each individual, generated by the proview command of MAPGD (version 0.4.26) (Ackerman et al. 2017). Individual-level disequilibrium (correlation of zygosity) was estimated with the methods of Lynch (2008) as implemented in Haubold et al. (2010). Results reported for each population are averages for each of the ten individuals with the highest sequence coverage, over a range of physical distances between sites of 1 and 250,000 bp. For the first 1,000 bp, estimates were obtained for increments of single nucleotides, whereas to offset the increased sampling variance with declining numbers of more distant pairs, for intersite distances of 1,000 to 20,000 bp, the data were binned into windows of 10-bp in width, and for larger distances into 100-bp bins.

### Analysis of selection on coding-region DNA

To infer the form of natural selection operating on the amino-acid sequences of protein-coding genes, we estimated nucleotide diversity at synonymous and nonsynonymous sites, *π_N_* and *π_S_*, for each gene within each population, as well as the ratio *π_N_ /π_S_*. For any particular site in a particular population, *π* is simply estimated as 2*p*(1 *− p*), where *p* is the minor-allele frequency estimate. To reduce sampling-variance problems associated with the generation of extreme values with ratios, we have opted to compute genespecific *π_N_ /π_S_* using the mean estimates of *π_N_* and *π_S_* across yearly samples to determine the ratios, as opposed to taking the mean *π_N_ /π_S_* across samples. The sampling variances of genespecific mean *π_N_*and *π_S_* were obtained by dividing the variance of population-specific estimates by the number of populations, and the sampling variance of *π_N_*and *π_S_*was then obtained by use of the Delta-method equation for the variance of a ratio (A1.19b, in Lynch and Walsh 1998).

To ascertain levels of between-species divergence, we estimated for each protein-coding gene *d_N_* and *d_S_,* respectively the mean numbers of nucleotide substitutions per nonsynonymous and synonymous sites, by drawing comparisons with orthologous genes contained within the closely related outgroup species *Daphnia obtusa.* We mapped sequence reads of *D. obtusa* to the KAP4 genome assembly (NCBI accession number GCF 021134715.1) using BWA-MEM (version 0.7.17) (https://arxiv.org/abs/1303.3997) and called genotypes using HGC (Maruki and Lynch 2017), setting the minimum and maximum coverage cut-off values at 6 and 100, respectively, and requiring the error-rate estimate *≤* 0.01. To avoid analyzing sites with mismapped nucleotide reads, we excluded sites involved in putatively repetitive regions identified by RepeatMasker. Codons containing undetermined nucleotides, gaps, and more than one nucleotide difference between species, and stop codons were removed from the alignments. *d_N_* and *d_S_* estimation was then restricted to genes with alignments consisting of at least 300 sites, using an algorithm that calculates potential numbers of replacement and silent sites (Hedrick 2005) in units of codons, weighted by allele frequencies in the *D. pulex* population, yielding divergence estimates similar to those by PAML (Yang 2007). For each gene, *d_N_* and *d_S_* were estimated as the mean divergence estimates across all populations, applying the Jukes and Cantor (1969) method. Then, genespecific *d_N_ /d_S_* estimates were obtained as the ratio of mean *d_N_*and *d_S_*estimates over years, and the variance of the ratio was again calculated using the Delta method.

### Enrichment analysis of gene ontology terms in the outlier genes

To infer the functions of outlier genes, we identified gene ontology (GO) terms (Ashburner et al. 2000) associated with each of the genes in KAP4 using Blast2GO (Götz et al. 2008). To find GO terms enriched in each type of outlier genes, we carried out enrichment analysis of the GO terms by a *χ*^2^ test using the Perl module Statistics::ChisqIndep (https://metacpan.org/pod/Statistics::ChisqIndep). To reduce the redundancy among GO terms enriched in outlier genes, we removed GO terms when at least 80% of the associated genes overlapped, using http://revigo.irb.hr/ .

### Computer simulations of fluctuating selection

To determine the effects of fluctuating selection on the long-term average within-population diversity, we employed the computational machinery described in Lynch (2020). This is a straight-forward “Wright-Fisher” population-genetic framework with discrete generations, fixed absolute population sizes, and biallelic loci.

Each generation involves consecutive episodes of mutation, selection, and random genetic drift. Mutation is bidirectional, and selection operates in an additive manner. Here, the simulations involve a single pair of completely linked loci, one neutral and the other subject to Gaussian fluctuations in the selection intensity on both of the alleles at the selected site each generation, with mean *s*, variance *σ*^2^, and no temporal correlation. The simulations are run long enough, after an ample burn-in period, to obtain stable estimates of the mean heterozygosity at each locus and of the long-term steady-state site-frequency spectra for both types of site. The code, written in C++, and is configured so that multiple populations with different sizes and/or mutation rates can be simultaneously run in parallel. The program used for these simulations, FlucSel22.cpp, is available at https://github.com/LynchLab/Fluctuating-Selection.

## Data availability

The FASTQ files of the raw sequencing data are available at the NCBI Sequence Read Archive (accession numbers PRJNA684968). The *D. pulex* genome assembly KAP4 is available at GenBank under accession GCF 021134715.1. The FASTQ files of the *D. obtusa* sequence reads are available at the NCBI SRA (accession number SAMN12816670), and the *D. obtusa* genome assembly is available at the NCBI GenBank (accession number JAACYE000000000).

## Acknowledgments

This work was supported by NIH grants R01-GM101672 and R35-GM122566-01 and NSF grant DEB-1257806 to ML. We thank Ken Spitze and Emily Williams for help in sample collection and DNA preparation.

## Notes

### Competing Interest Statement

The authors have declared no competing interest.

