## Supplementary Figures for "Evolutionary Insights from a Large-scale Survey of Population-genomic Variation"

**Supplemental Figure S1**. Chromosome distribution of genes with *π_n_/π_s_* significantly *>* 1*.*0. Red bars indicated centromere regions.

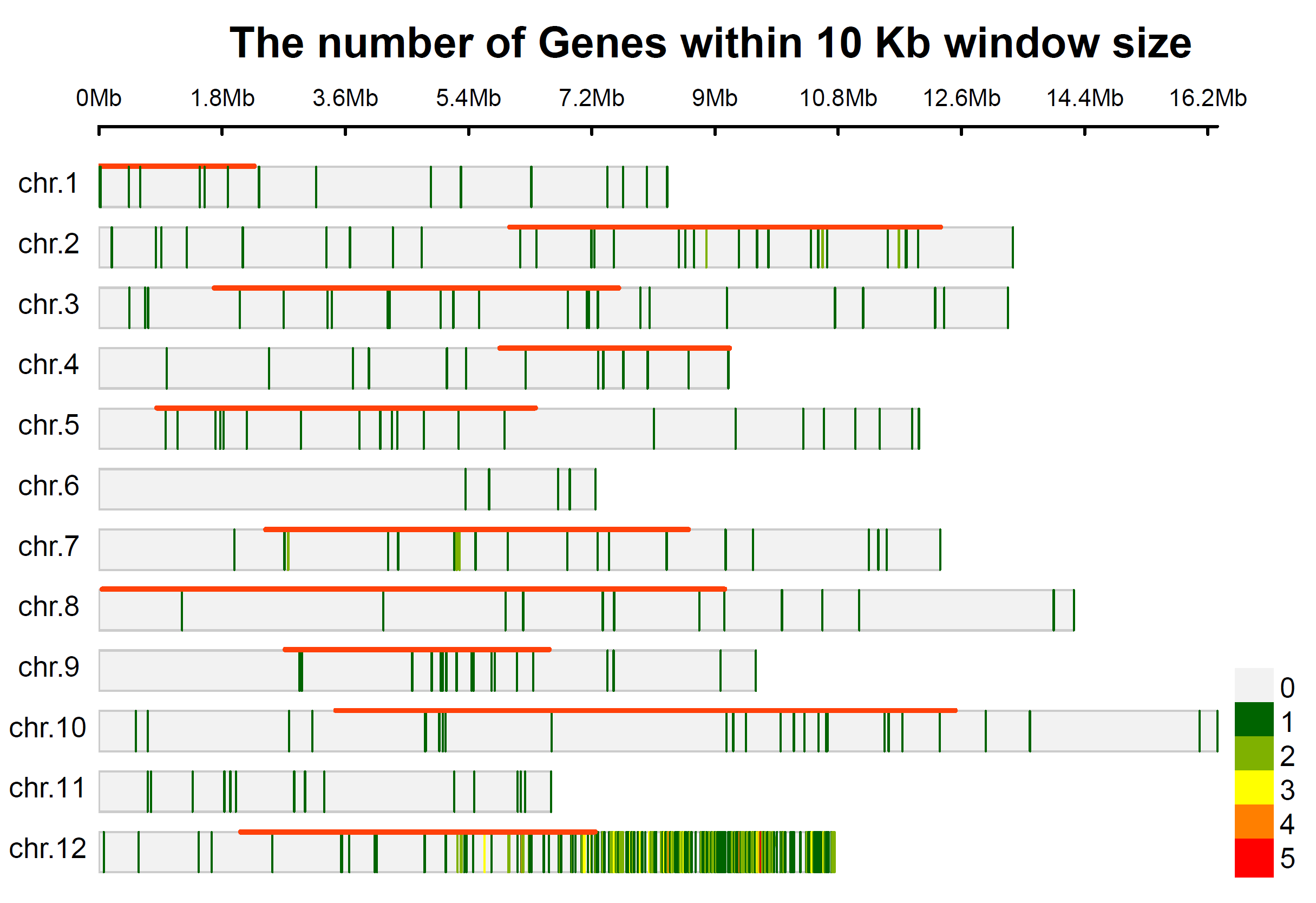

**Supplemental Figure S2**. Chromosome distribution of genes with NI significantly < 1*.*0. Red bars indicated centromere regions.

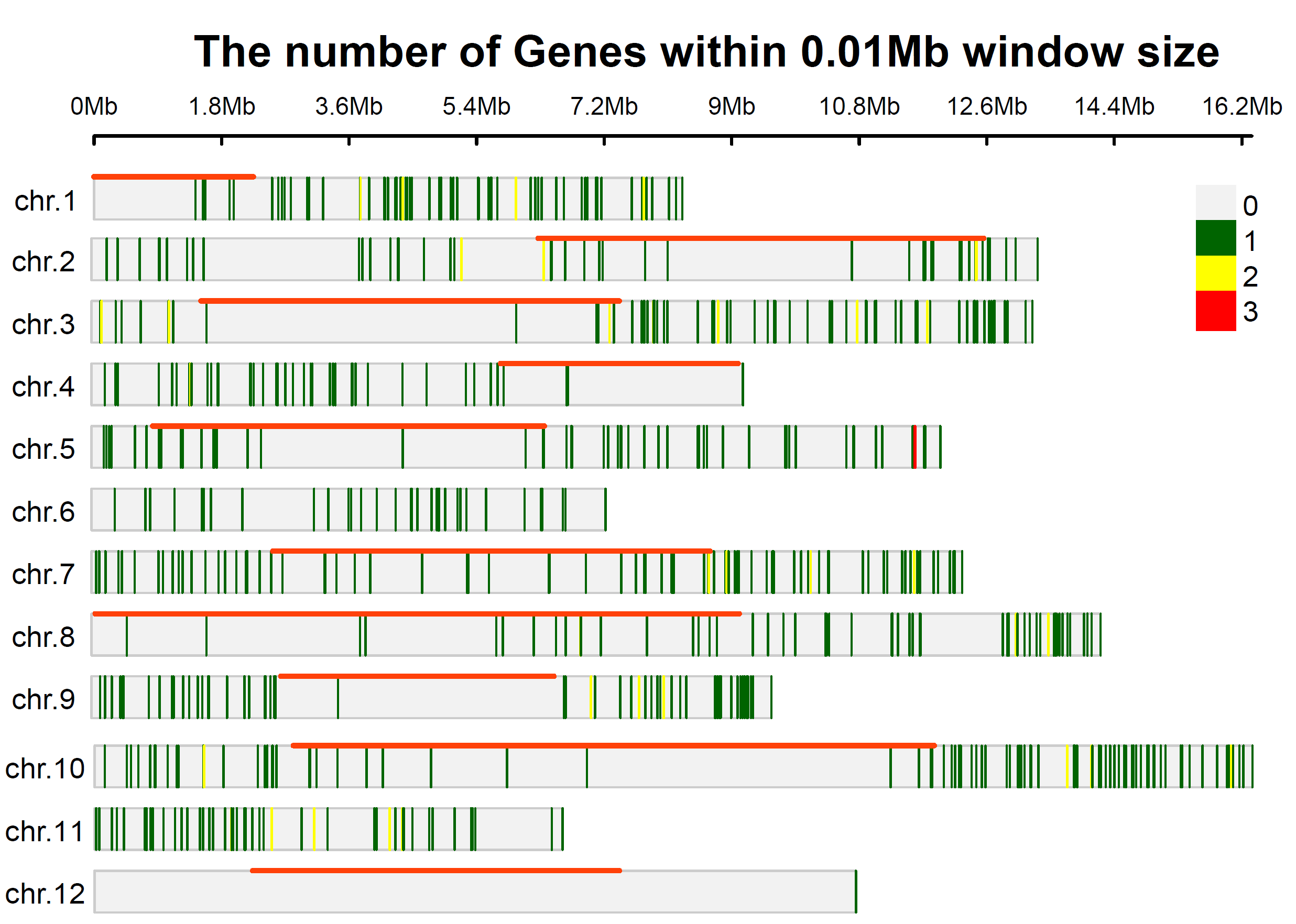
